## supplemental figures 1-9 for "Virtual Mouse Brain Histology from Multi-contrast MRI via Deep Learning"

### Supplementary Materials

### Supplementary Fig. 1

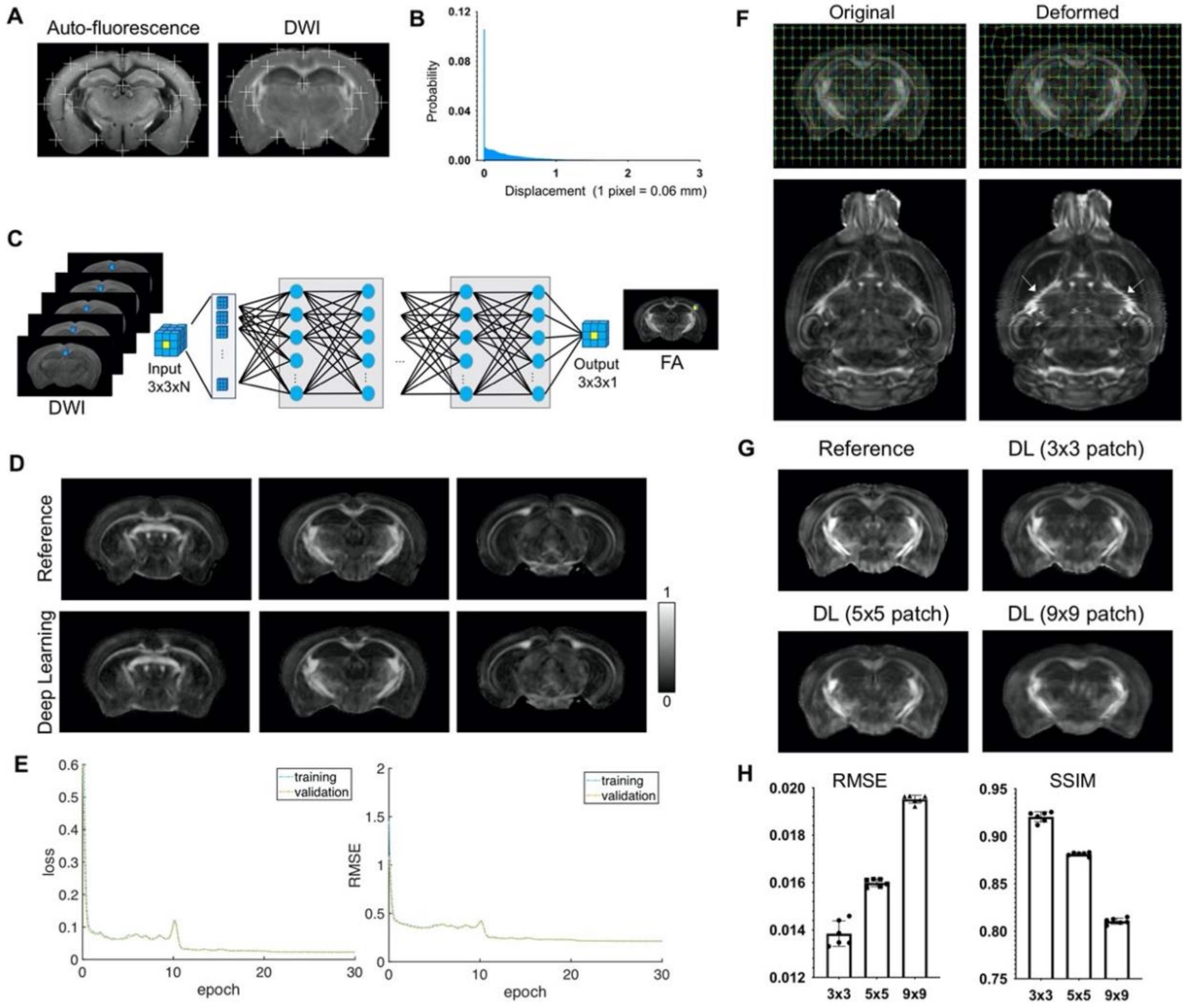


**Supplementary Fig. 1:** Evaluate the effects of mismatches between input MRI data and target auto-fluorescence (AF) data on deep learning outcomes **A:** The overall registration accuracy is visually examined by overlaying a set of landmarks on AF and average diffusion-weighted (DWI) images. **B:** Distribution of pixel displacement due to mismatches between AF and MRI data were estimated using image mapping. Overall, 70% of the pixel displacements are within 1 pixel (0.0625 mm) and 95% within 2 pixels (0.125mm). **C:** A convoluted neural network with a similar archittecture as the one in the main text was trained using DWIs of *ex vivo* mouse brains as inputs and corresponding maps of fractional anisotropy (FA), generated by fitting the DWIs to a diffusion tensor model, as targets. In this case, the inputs and targets are perfectly co-registered. **D:** Comparisons of FA maps generated from model fitting and from the convolutional neural network (Deep Learning). Overall, the deep learning results show good agreement with the reference from perfectly registered input and target data. The 3x3 patch size used by the network caused smoothing in the deep learning results. **E:** Smoothed curves of mean square error loss (left) and RMSE (right) with respect to the reference FA maps during training measured on the training (blue) and validation (yellow) datasets. Each epoch is 300 iterations. **F:** Two-dimensional random displacement fields with the same distribution as shown in **A** were introduced to deform FA maps in **C**. The white arrows in the horizontal images indicated the misalignments introduced by this method compared to the original FA maps. Notice the zip-zagged boundaries in the deformed FA map compared to the smooth boundaries in the original FA map. **G:** Deep learning results generated using different patch sizes. Larger patch size will be able to accommodate more mismatches between input and target data but also increase image smoothing in the results. For the amounts of residual mismatches shown in **A**, the 3x3 patch size is robust to the mismatches with minimal smoothing effects. **H:** RMSE and SSIM values of results generated using different patch sizes. There are significant differences between different patch sizes (t-test, p<0.00001)

**Supplementary Fig. 2**

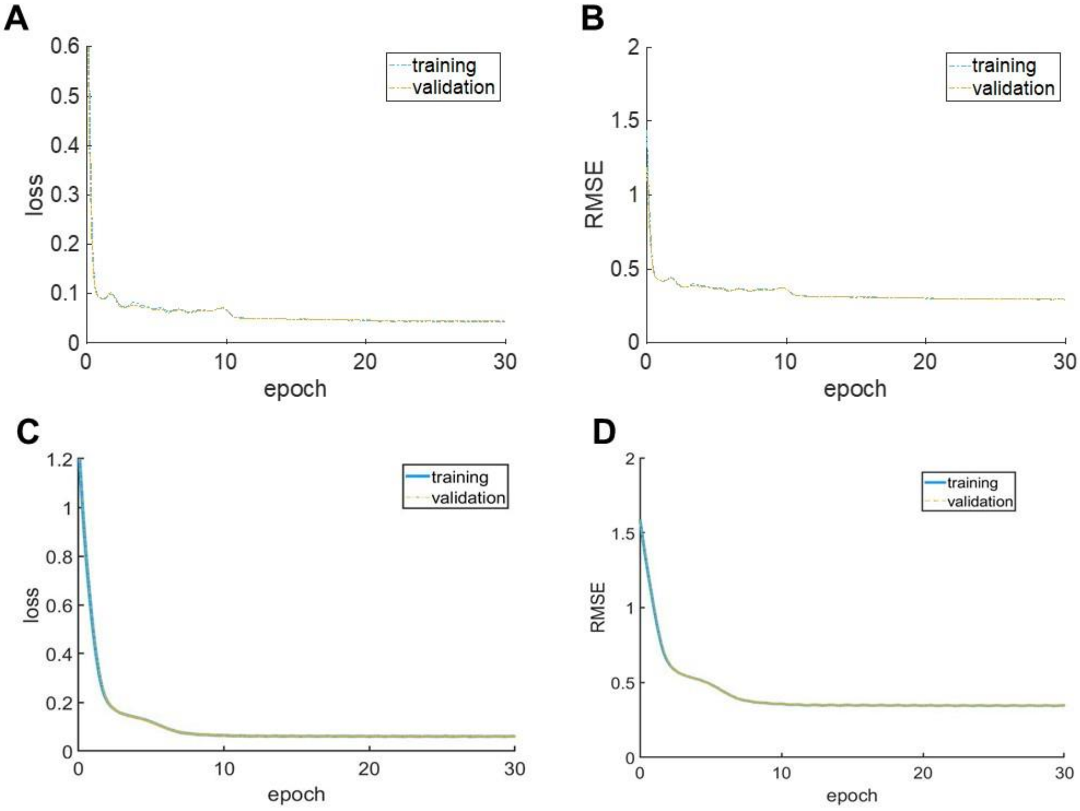


**Supplementary Fig. 2:** Training convergence curves of MRH-AF network (A-B) and MRH-MBP during transfer learning. Smoothed curves of mean square error loss (A, C) and RMSE (B, D) with respect to the reference during training measured on the training (blue) and validation (yellow) datasets. Each epoch is 300 iterations.

**Supplementary Fig. 3**

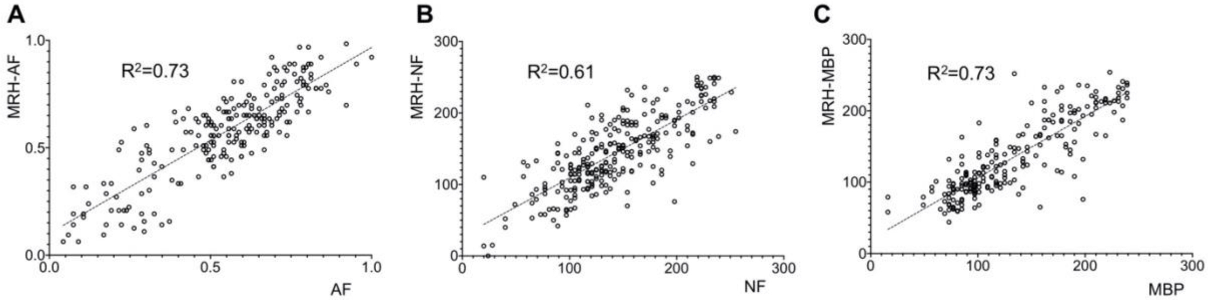


**Supplementary Fig. 3:** Strong correlations between MRH generated results based on test MRI data and reference data for auto-fluorescence (AF), neurofilament (NF), and myelin basic protein (MBP). P values for all these tests are less than 0.001.

**Supplementary Fig. 4**

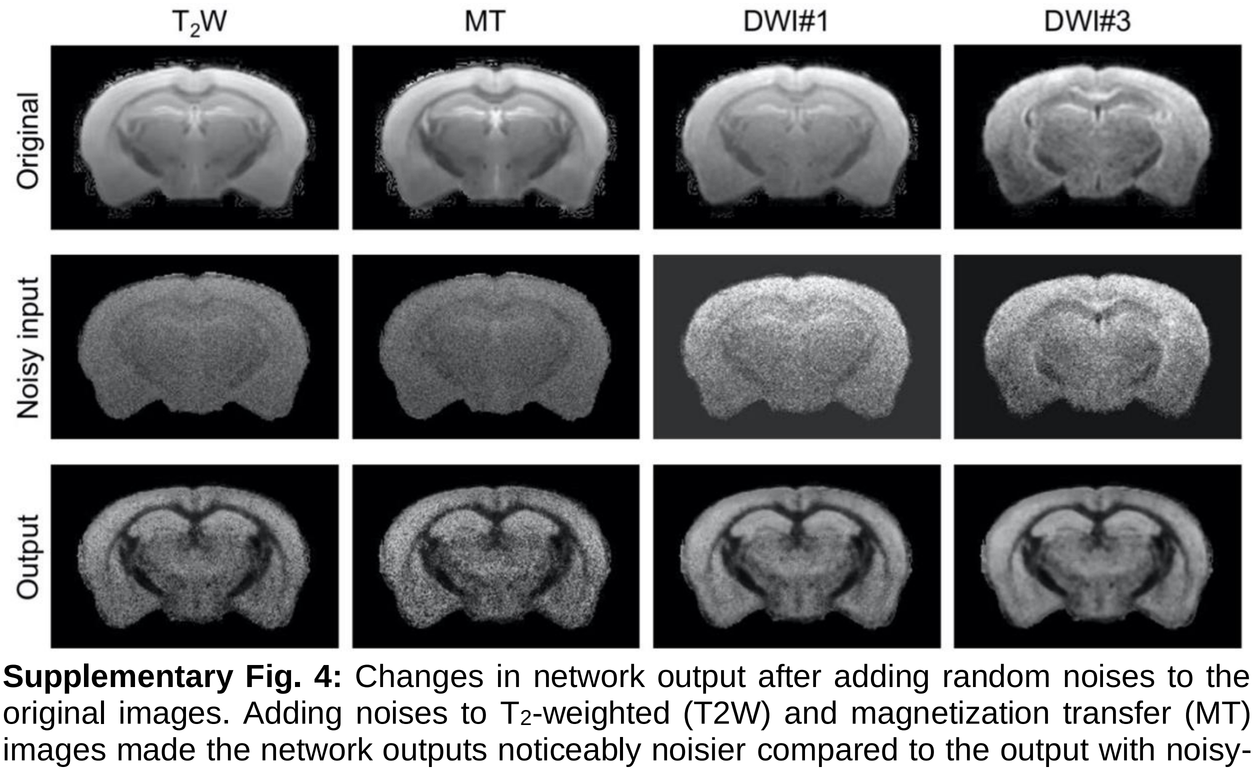


**Supplementary Fig. 4:** Changes in network output after adding random noises to the original images. Adding noises to $T_2$-weighted (T2W) and magnetization transfer (MT) images made the network outputs noticeably noisier compared to the output with noisy-free inputs. In comparison, similar level of noises added to two diffusion weighted images (DWI#1 and DWI#3) produced less apparent changes in the output.

**Supplementary Fig. 5**

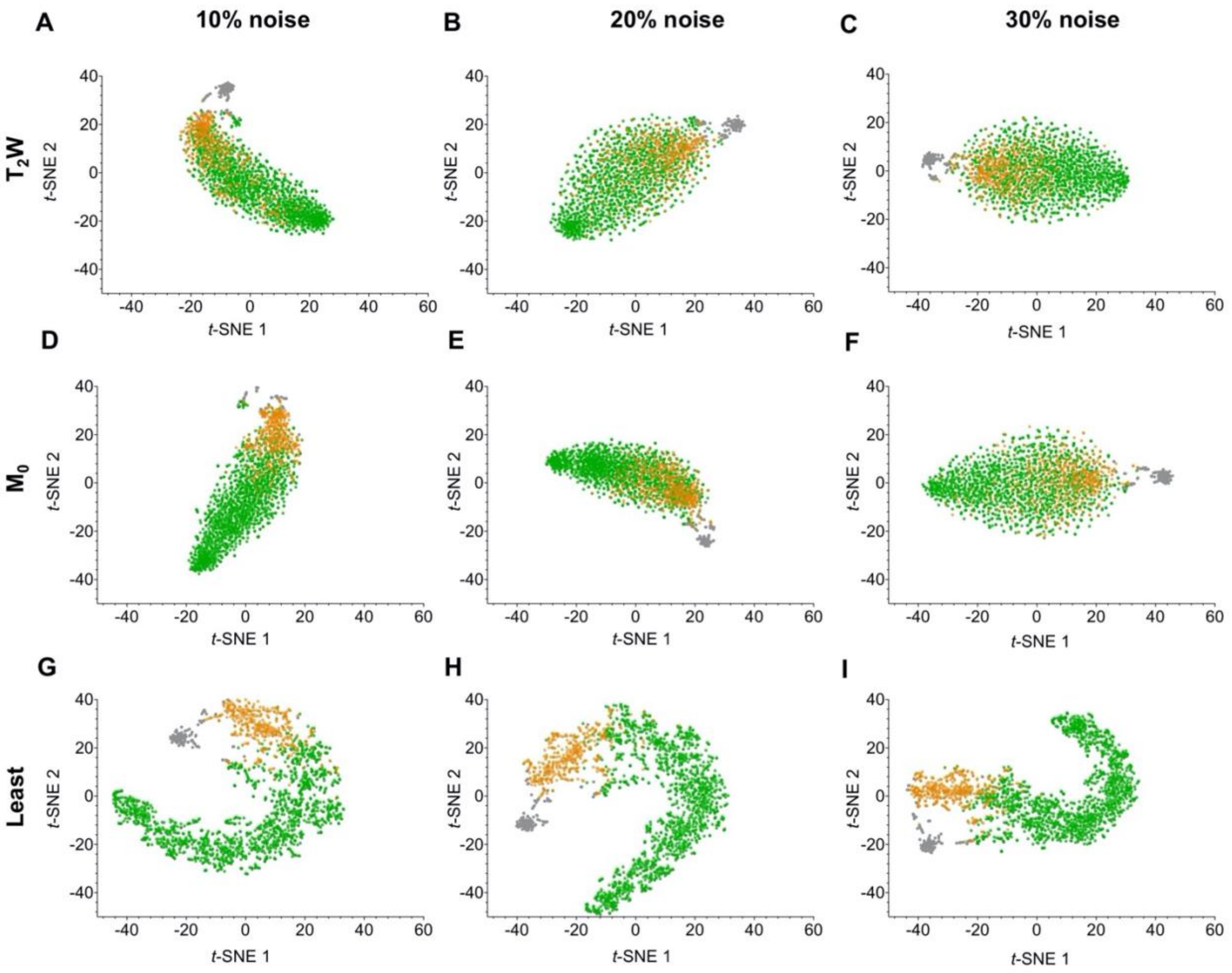


**Supplementary Fig. 5:** The effects of adding random noises to images with high and low contributions in MRH-AF. Adding 10%, 20%, and 30% noises to $T_2$-weighted ($T_2$W) and baseline MT ($M_0$) images, which have the highest contributions in MRH-AF, results in less well separated clusters in the feature space defined by *t*-SNE analysis compared to the noise-free result shown in Fig. 1G, whereas adding similar levels of noises to the MR image with the least contribution results in less apparent changes in the *t*-SNE result.

**Supplementary Fig. 6**

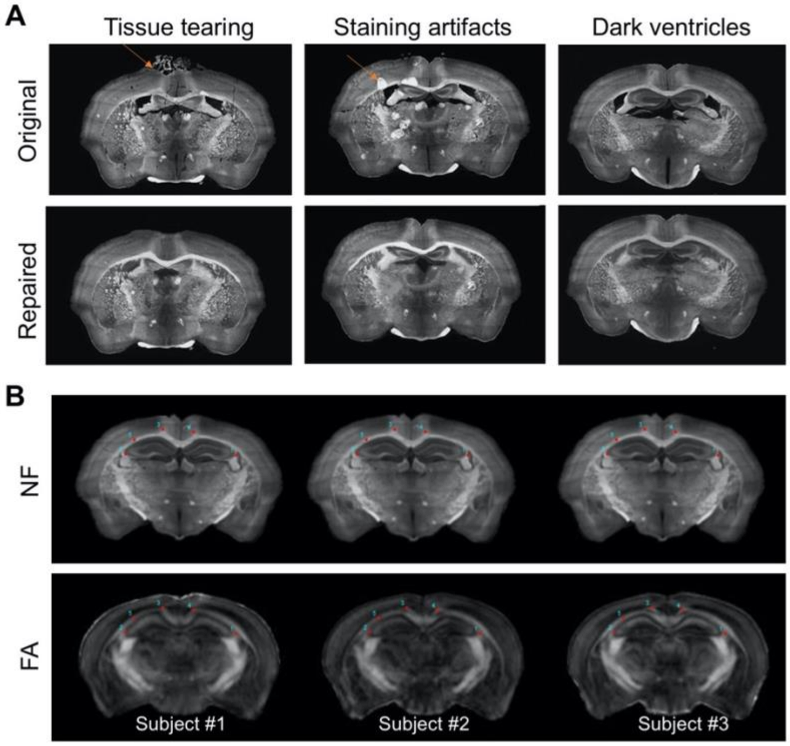


**Supplementary Fig. 6:** Preparation and co-registration of serial 2D histological sections to MRI data. **A:** Two types of common artifacts in neurofilament (NF)-stained images were repaired, and the ventricular spaces were filled with the average intensity values of the cortex. **B:** Examples of co-registered histological and MRI data. Both NF-stained histological images and ex vivo MRI data were aligned to the space defined by ARA. Manually placed landmarks were overlaid to show the overall registration quality. The fractional anisotropy (FA) maps are shown here because white matter structures usually have high FA values due to the coherently arranged axons.

**Supplementary Fig. 7**

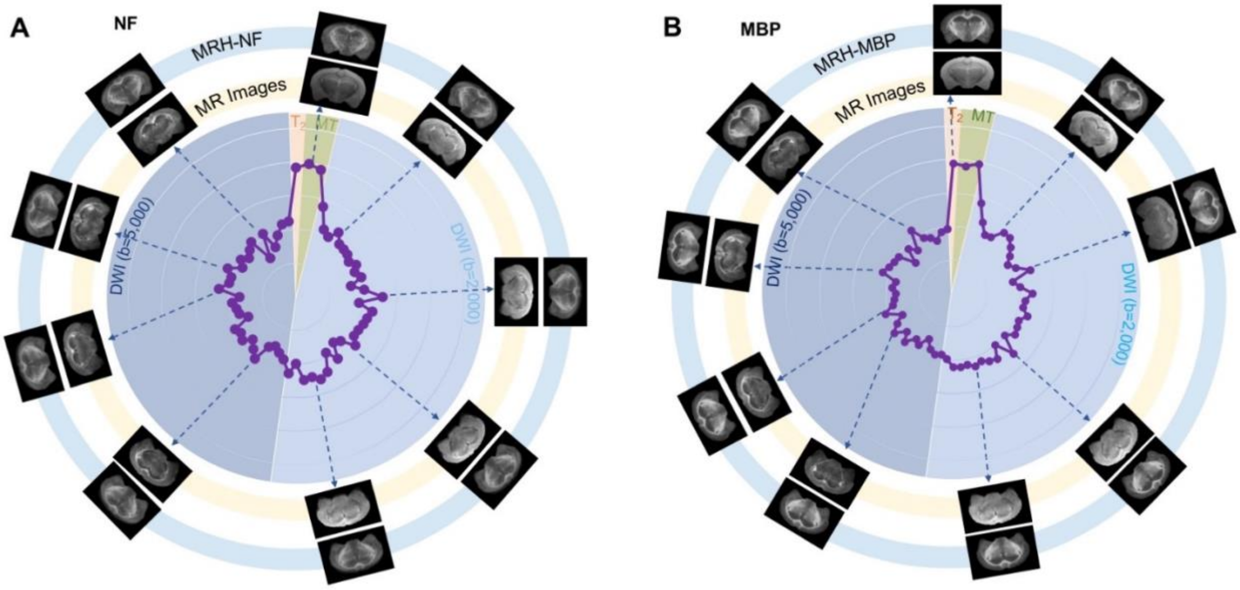


**Supplementary Fig. 7:** Plots of the contributions of 67 MR images in MRH-NF (A) and MRH-MBP (B). $T_2$-weighted ($T_2$) and magnetization transfer (MT) images show the highest contributions in both cases. In both plots, the contributions are normalized by the total contribution of all MR images. Images displayed on the outer ring (light blue, MRH-NF/MBP) show the network outcomes after adding 10% random noises to a specific MR image on the inner ring (light yellow).

**Supplementary Fig. 8**

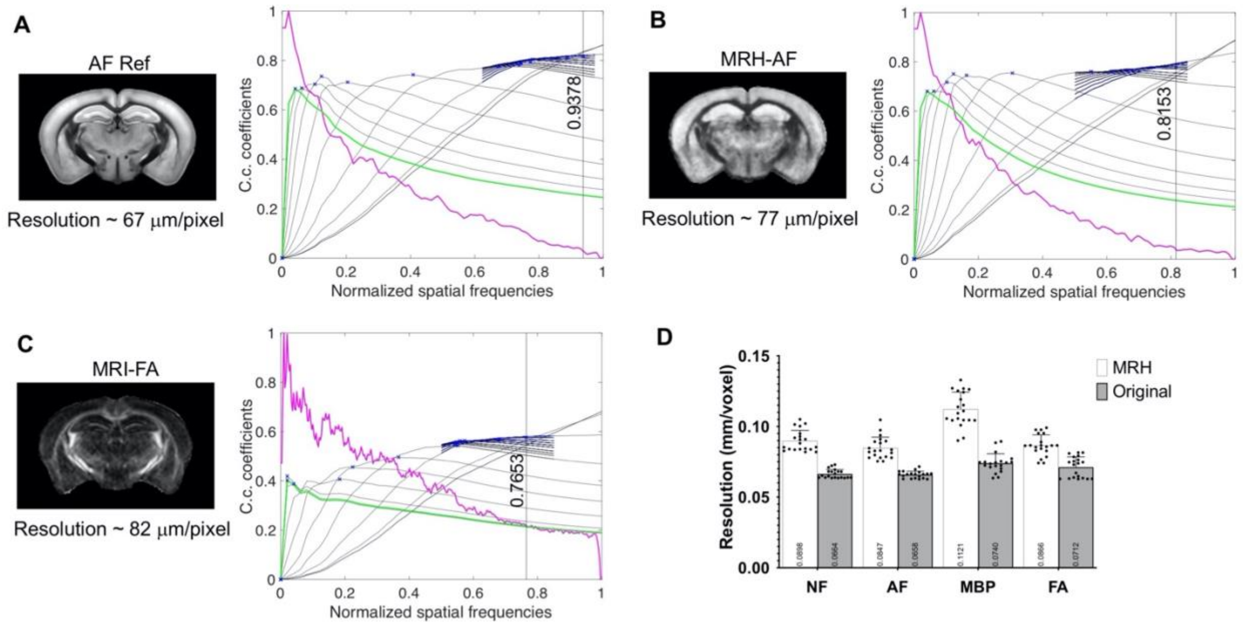


**Supplementary Fig. 8:** Resolution of down-sampled AF reference data, MRH-AF data, and FA map of the input MRI data estimated using deconvolution analysis. The top row shows the actual images and the bottom row shows the deconvolution analysis results. The estimated resolutions were computed by dividing the nominal resolution (62.5 um/pixel) by the highest normalized spatial frequency detected in the images (indicated by the vertical lines). **D:** The resolutions of input MRI and histological data were higher than the resolutions of MRH outputs based on the deconvolution analysis (paired t-test, $p<0.000001$). In this case, the histological data had been first down sampled to 0.1 mm/pixel.
